# Revisiting *Cryptococcus* extracellular vesicles properties and their use as vaccine platforms

**DOI:** 10.1101/2020.08.17.253716

**Authors:** Juliana Rizzo, Sarah Sze Wah Wong, Anastasia D. Gazi, Frédérique Moyrand, Thibault Chaze, Pierre-Henri Commere, Sophie Novault, Mariette Matondo, Gerard Pehau-Arnaudet, Flavia C. G. Reis, Matthijn Vos, Lysangela R Alves, Robin C. May, Leonardo Nimrichter, Marcio L. Rodrigues, Vishukumar Aimanianda, Guilhem Janbon

## Abstract

Whereas extracellular vesicle (EV) research has become commonplace in different biomedical fields, this field of research is still in its infancy in mycology. Here we provide a robust set of data regarding the structural and compositional aspects of EVs isolated from the fungal pathogenic species *Cryptococcus neoformans, C. deneoformans and C. deuterogattii*. Using cutting-edge methodological approaches including cryogenic electron microscopy and cryogenic electron tomography, proteomics, and flow cytometry, we revisited cryptococcal EV features and suggest a new EV structural model, in which the vesicular lipid bilayer is covered by mannoprotein-based fibrillar decoration, bearing the capsule polysaccharide as its outer layer. About 10% of the EV population is devoid of fibrillar decoration, adding another aspect to EV diversity. By analyzing EV protein cargo from the three species, we characterized the typical *Cryptococcus* EV proteome. It contains several membrane-bound protein families, including some Tsh proteins bearing a SUR7/PalI motif. The presence of known protective antigens on the surface of *Cryptococcus* EVs, resembling the morphology of encapsulated virus structures, suggested their potential as a vaccine. Indeed, mice immunized with EVs obtained from an acapsular *C. neoformans* mutant strain rendered a strong antibody response in mice and significantly prolonged their survival upon *C. neoformans* infection.

## 1. Introduction

All living organisms release lipid bilayer-delimited particles defined as extracellular vesicles (EVs) (Deatherage and Cookson 2012, Witwer and Théry 2019). EV size ranges from 20 nm to close to one micrometer in diameter. In mammalian cells, two major classes of EVs have been defined, microvesicles and exosomes, according to their size and cellular origin (Meldolesi 2018, van Niel et al. 2018). In these organisms, a large body of literature describes how EVs participate in intercellular signaling within and in organism-to-organism communication, including carcinogenesis and host-pathogen interactions (Xu et al. 2018, Shopova et al. 2020). In fungi, the first report of fungal EVs was published in 2007 (Rodrigues et al. 2007), and, since then, their existence has been reported in many species of pathogenic and nonpathogenic fungi (Rizzo et al. 2020).

By analogy with mammalian EVs, it has been hypothesized that fungal EVs could also participate in many biological processes (Rodrigues and Casadevall 2018). Indeed, some reports indicate their relevance in diverse mechanisms related to fungal pathophysiology, such as antifungal resistance and biofilm formation (Leone et al. 2018, Zarnowski et al. 2018), transfer of virulence-associated molecules and modulation of host cells (Oliveira et al. 2010, Vargas et al. 2015, Rizzo et al. 2017, Bielska et al. 2018, Souza et al. 2019, Hai et al. 2020), cell wall remodeling and biogenesis (Zhao et al. 2019, Rizzo et al. 2020), among others (Bielska and May 2019, Rizzo et al. 2020). Nevertheless, the molecular mechanisms implicated in these exchanges of information, as well as the genetics regulating fungal EV biogenesis and release, remain elusive.

As with their mammalian, bacterial and plant counterparts, fungal EVs have been shown to contain proteins, pigments, nucleic acids, polysaccharides, lipids, among other molecules (Eisenman et al. 2009, Vallejo et al. 2012, da Silva et al. 2015, Rodrigues et al. 2015, Joffe et al. 2016, Rayner et al. 2017, Reis et al. 2021). Besides the claudin-like Sur7 family proteins, recently suggested as EV protein markers in *Candida albicans* (Dawson et al. 2020), no other fungal EV specific molecular marker has been reported. Indeed, the laborious and inefficient EV isolation protocols that have been used until recently have strongly limited the knowledge on their composition. Additional hurdles regarding purification methods potentially affect an accurate vesicular compositional characterization (Théry et al. 2018), including potential carryover contaminants such as protein aggregates (Chiou et al. 2018). Regarding EV morphological diversity, previous studies have reported the heterogeneity of fungal EV size, as recently reviewed (Bielska and May 2019). However, single particle analyzers such as the widely used Nanoparticle Tracking Analysis (NTA) and most common flow cytometers cannot reliably evaluate particles smaller than 100 nm in diameter (Maas et al. 2015, Théry et al. 2018, Chiang and Chen 2019, Noble et al. 2020). Overall, although a considerable number of fungal EV-related studies have been published in recent years, our knowledge of fungal EV structure and composition remains limited, which prevents further robust analysis of their functions.

Seven species of pathogenic *Cryptococcus* have been described (Hagen et al. 2015). Whereas species belonging to the *neoformans* clade *(C. neoformans* and *C. deneoformans)* typically affect immunocompromised patients, species belonging to the *gattii* clade (*C. gattii*, *C. deuterogattii*, *C. tetragattii*, *C. decagattii*, and *C. bacillisporus)* are often primary pathogens and can cause aggressive pulmonary infections as well as meningoencephalitis (Kwon-Chung et al. 2014, Rajasingham et al. 2017, Janbon et al. 2019). *C. neoformans* has historically been one of the most studied fungi regarding EV biology (Rodrigues et al. 2007, Bielska and May 2019, Rizzo et al. 2020). However, the only structural analyses of EVs from this organism are now very outdated and technologies used have shown tremendous improvements since then (Emelyanov et al. 2020, Noble et al. 2020).

To date, *Cryptococcus* EV proteomic approaches have identified 92 and 202 proteins, with very poor overlap and no evaluation of their abundance or enrichment (Rodrigues et al. 2008, Wolf et al. 2014). Finally, although the current model of fungal EV structure contains proteins located on the vesicular surface (by analogy with the mammalian EV structures (Emelyanov et al. 2020, Noble et al. 2020)), more experimental evidence is necessary to identify these putative membrane-associated molecules. Since many immunogenic proteins are often found to be associated with EVs, their vaccine potential has been explored mostly for bacterial and parasitic infections (Coakley et al. 2017, Wang et al. 2018), and more recently also for fungal infections (Colombo et al. 2019, Vargas et al. 2020).

In the present study, we used the recently described protocol (Reis et al. 2019) to obtain EV-enriched samples from *Cryptococcus*, together with cutting edge methodological approaches to revisit *Cryptococcus* EV structure, cargo, and their biological functions. Here we report a detailed analysis of three species (*C. neoformans*, *C. neoformans* and *C. deuterogattii*) for which both a good genome annotation and RNA-Seq data were available (Janbon et al. 2014, Gonzalez-Hilarion et al. 2016, Gröhs Ferrareze et al. 2021). We produced a robust set of data containing cryo-electron microscopy (cryo-EM) and cryo-electron tomography (cryo-ET) proteomics, and nanoscale flow cytometry analysis, suggesting a new EV structural model, in addition to a list of cryptococcal EV protein markers. Our results led us to further evaluate the EV biological roles in murine models, emphasizing their potential use as an anti-cryptococcosis vaccine.

## 2. Material and Methods

### Strains and media

The wild type strains used in the study were *C. neoformans* strain KN99α, *C. deneoformans* strain JEC21, *C. deuterogattii* strain R265, *C. albicans* strain SC5314, and *S. cerevisiae* strain S288C. The *C. neoformans* strain NE367 (*MAT*α *cap59Δ::NAT)* has been previously described (Moyrand et al. 2007). The strains *MAT*α *vep1Δ::NAT* (CNAG_03223), *MAT*α *hoc3Δ::NAT* (CNAG_00158), *MAT*α *alg3Δ::NAT* (CNAG_05142), *MAT*α *ktr3Δ::NAT* (CNAG_03832) have been constructed in the Hiten Madhani lab (UCSF, USA) and obtained from the Fungal Genetic Stock Center. To construct the strains NE1281 (*MAT*α *mp88Δ::NEO*) and NE1469 (*MAT*α *vep1Δ::NAT mp88Δ::NEO*), we replaced the entire CNAG_00776 (*MP88*) CDS by the NEO marker in the strains KN99α and *MAT*α *vep1Δ::NAT*, respectively. We here followed the previously described CRISPR CAS9 method (Fan and Lin 2018). The plasmid pPZP-NEO1 used to amplify the *NEO* selective marker was kindly provided by Dr. Joseph Heitman (Duke University). The deletion cassettes was constructed using a strategy previously applied to *Neurospora crassa* (Collopy et al. 2010). The transformants were then screened for homologous integration, as previously described (Moyrand et al. 2007). Two representatives independently obtained mutant strains were stocked at −80°C. All primer sequences used are provided in Table S1. All strains were taken from the G. Janbon laboratory collection at −80°C, plated on yeast extract-peptone-dextrose (YPD) and incubated at 30°C for 48h before each experiment

### EV isolation, labelling and proteinase K treatment

EV purification was based on the recently published protocol (Reis et al. 2019) with some modifications. One loop of cells was inoculated into 10 mL of liquid YPD and incubated at 30°C for 24h with shaking (150 rpm). Cells were washed twice with 10 ml of sterile water, counted and diluted to a density of 3.5 x 10^7^ cells/mL in water. Aliquots of 300 µL of the cellular suspension were spread onto synthetic dextrose (SD) solid medium plates and incubated for 24 h at 30°C to reach cell confluence. The cells were recovered from each plate with a 10 µL inoculation loop and transferred to an ultracentrifugation tube containing 10 mL 0.22 µm-filter sterile of 0.01 M PBS. Cells were homogenized and collected by centrifugation at 5,000 x g for 15 min at 4☐C. Supernatants were collected and centrifuged again at 15,000 x g for 15 min at 4☐C to eliminate cellular debris. The resting supernatants were filtered through 0.45µm syringe filters and ultracentrifuged at 100,000 x g for 1h at 4°C (SW41 Ti swinging-bucket rotor, Beckman Coulter). The supernatant was discarded and the pellet suspended in 0.22 µm-pore filtered or 0.02 µm-pore filtered (for Flow Cytometry analysis) PBS for immediately use or stored at −80°C for further experiments. The amount of total sterol in the EV samples was measured by the Amplex™ Red Cholesterol Assay Kit (ThermoFisher, A12216) and adjusted for the subsequent experiments.

EVs were labelled either with the Concanavalin A (ConA) - Alexa Fluor™ 488 conjugated, or with the Alexa 488 labelled anti-GXM monoclonal antibody 18B7 (Casadevall et al. 1992), a kind gift of Oscar Zaragoza. The ConA stock solution (5mg/mL) was previously centrifuged at 13.000 x rpm for 2 min, in order to eliminate possible aggregates, and diluted to 500 µg/mL in filtered PBS. In 1.5 mL Eppendorf tubes, 5 µL of ConA (500 µg/mL), together with 5 µL EV suspension were add to a final volume of 100 µL filtered PBS. The tubes were incubated for 1 h at 30°C, under agitation and protected from light. After incubation, 10 mL of 0.02 µm-pore filtered PBS were added to the EV suspension and then submitted to ultracentrifugation for 1 h at 100 000 x g at 4°C. The supernatant was again discarded and pellets suspended in 300 µL of 0.02 µm-pore filtered before being transferred to BD Trucount™ Tubes (BD Biosciences) and proceeded to Flow Cytometry analysis. A similar protocol was applied for the EV labelling with the Alexa 488 labelled anti-GXM monoclonal antibody 18B7, which was diluted 20 times before adding to EV suspension.

EV proteinase K treatment was performed following the previously described protocol (Yang et al. 2021) with some modifications. Briefly, proteinase K was added to the EV suspension (0.17 µg of sterol) to a final concentration of 2 mg/mL in 0.02 µm-pore filtered PBS. After proteolysis for 1h at 55°C under agitation (300 rpm), the enzymatic reaction was stopped by the proteinase inhibitor PMSF (1 mM) for 20 min at RT. Proteinase K treated EVs were finally submitted to ConA labelling, ultracentrifuge washed as described before and analyzed by flow cytometry. Control conditions included untreated EVs and EVs incubated only with PMSF.

### Flow cytometry

EVs were analyzed and sorted on a cell sorter MoFlo Astrios (Beckman Coulter) equipped with an EQ module specifically developed to detect nanoparticles and with 488 nm and 561 nm lasers at 200 mW. The sorting was carried out with a 70 μm nozzle at a pressure of 60 PSI and a differential pressure with the sample of 0.3-0.4 PSI. The sheath liquid NaCl 0.9% (REVOL, France) was filtered on a 0.04 μm filter. The analyses were on the SSC parameter of laser 561, with threshold set to 0.012% in order to have maximum 300 eps. An M2 mask was added in front of the FSC. All SSC and FSC parameters are viewed in logarithmic mode. The calibration of the machine was carried out using Megamix-Plus SSC beads from BioCytex. We used the Trucount™ Tubes to normalize the EV counting for ConA labelling, and the fluorescence of the Mab18B7 and alexa 488 conjugated, and beads Trucount™ was measured on parameter 488-513/26. Control conditions including ultracentrifuge washed PBS, previously incubated with ConA were used to evaluate the PBS associated noise and to normalize labelling percentages. Flow Cytometry data were analyzed by FlowJo V10 Software.

### Nanoparticle tracking analysis (NTA)

Quantitative determination of EV size distribution was performed by NTA, in addition to microscopic methods. Protocols that were recently established for the analysis of cryptococcal EVs were used (Reis et al. 2019). Briefly, ultracentrifugated pellets were 20- to 50-fold diluted in filtered PBS and measured within the optimal dilution range of 9 x 10^7^ to 2.9 x 10^9^ particles/mL on an LM10 nanoparticle analysis system, coupled with a 488-nm laser and equipped with an SCMOS camera and a syringe pump (Malvern Panalytical, Malvern, United Kingdom). The data were acquired and analyzed using the NTA 3.0 software (Malvern Panalytical).

### Cryo-EM and cryo-ET

EVs (4µL) were spotted on glow-discharged lacey grids (S166-3, EMS) and cryo-fixed by plunge freezing at −180°C in liquid ethane using a Leica EMGP (Leica, Austria). Grids were observed either with Tecnai F20, or Titan Krios (Thermo Fisher Scientific). The Tecnai F20 (Thermo Fisher Scientific) was operating at 200kV and images were acquired under low-dose conditions using the software EPU (Thermo Fisher Scientific) and a direct detector Falcon II (Thermo Fisher Scientific).

Cryo-electron tomography was performed using 5 nm protein-A gold particles (UMC, Utrecht). These were mixed with the sample to serve as fiducial markers for subsequent image alignment. EV sample (4 μL) was applied to glow discharged Lacey grids (S166-3, EMS) prior plunge-freezing (EMGP, Leica). Initial bi-directional tilt series acquired using a TECNAI F20 transmission electron microscope (FEI) operated at 200kV under parallel beam conditions using a Gatan 626 side entry cryoholder. The SerialEM software (Mastronarde 2005, Schorb et al. 2019) was used to automatically acquire images every 2° over a ±45° range using a Falcon II direct detector with a pixel size of 2 Å, using a total dose of 180 electrons per Å2. At least 100 EV cryo-EM images obtained from TECNAI F20 were used for measuring EV diameter and decoration thickness in wild type (WT) and mutant strains. For each EV, an average of three different measurements were used to calculate the diameter (delimited by the lipid bilayer) and the decoration thickness.

Dose-symmetric tilt series were collected on a 300kV Titan Krios (Thermo Scientific) transmission electron microscope equipped with a Quantum LS imaging filter (Gatan, slit with 20☐eV), single-tilt axis holder and K3 direct electron detector (Gatan). Tilt series with an angular increment of 2° and an angular range of ±60° were acquired with the Tomography software (Thermo Scientific). The total electron dose was between 120 and 150 electrons per Å2 and the pixel size at 3.38☐Å. Dose symmetric tilt series were saved as separate stacks of frames and subsequently motion-corrected and re-stacked from −60° to +60° using IMOD’s function align frames (Mastronarde and Held 2017) with the help of a homemade bash script.

Initial image shifts were estimated using IMOD’s function tiltxcorr. Alignments were further optimized in IMOD using the tracing of 30-40 gold fiducials across the tilt series. The fiducial models gave an overall of a fiducial error around 6 ± 2.7 Å. In cases of a higher error, local alignments were taken into consideration, to further correct the sample’s beam induced motion observed. Three-dimensional reconstructions were calculated in IMOD by weighted back projection using the SIRT-like radial filter to enhance contrast and facilitate subsequent segmentation analysis.

### EV-modeling and analysis of tomographic data

Tomograms were displayed and analyzed using the 3dmod interface of IMOD (Kremer et al. 1996). EVs were modeled with manual tracing of their great circle prior the use of the spherical interpolator of IMOD. If the elliptical contours calculated could not follow the vesicular membrane adequately, further manual tracing was used before re-applying the interpolator. This involved tracing of membranes near the poles of the vesicles where the membrane information could still be followed. To evaluate and assign diameters to a total of 434 *C. neoformans* regular vesicles, located in 39 tomograms, the value of the perimeter of the spheroid’s great circle was extracted using the imodinfo function of IMOD, from the same initial manually traced contours used for modelling. To display in 3D the vesicle contour data were meshed using the imodmesh function of IMOD. The projections of the 3D spheroidal models were displayed and rotated to study their 3D geometry.

For the evaluation of the decoration thickness, regular vesicles were analyzed by manually measuring the outer EV diameter (delimited by the fibrillar decoration) and the inner diameter (delimited by the lipid bilayer), across the longest axis of the vesicle. The final calculation of the decoration thickness was the subtraction of the inner diameter from the outer diameter, divided by two. For the modeling of the fibrillar decoration, the IMOD surface models were imported to UCSF Chimera (Pettersen et al. 2004). The models were used as masks to extract a slab of data around their outer surface, corresponding to the decoration. The thickness of the slab used refers to the mean value provided by the aforementioned manual analysis. Iso-surface representation of the decoration and final 3D data visualization of the models performed with UCSF Chimera (Pettersen et al. 2004).

### Immunization assays

The animal experiments were approved by the ethical committee for animal experimentation Comité d’Éthique en Experimentation Animale (CETEA Project license number 2013-0055). Six-week old female BALB/c mice (Janvier Labs) were used for immunization study. The amount of EVs, in protein concentration, was determined by BCA method prior to immunization. Following, three intraperitoneal injections (fixing protein concentration in the EVs to either 1 or 10 µg and suspending in 100 μL PBS) at 15-day intervals were given to the mice. The control group of mice was injected only with PBS. Blood was collected from the submandibular veins of the mice three days after the last immunization and just before the fungal infection and tested for antibody response by Western blot. Briefly, the EVs-associated proteins were separated on 12% SDS-PAGE, and electroblotted to nitrocellulose membrane. By Western blotting, using the mouse sera at dilution 1:1000 and anti-mouse IgG antibody conjugated to peroxidase (Sigma Aldrich), the antibody response specific to the EV-associated proteins was examined. Once the antibody response was confirmed, all the immunized and control mice were challenged intranasally, around one month from the last immunization, with 1 x 10^4^ cells of *C. neoformans* wild-type strain, and their body weights and survival were monitored until all mice succumbed to the infection. The immunization assay was performed in two biological replicates.

### Vesicle denaturation and protein digestion

EVs proteins were solubilized in urea 8 M, Tris 100 mM pH 7.5, 5 mM tris (2-carboxyethyl) phosphine (TCEP) for 20 min at 23°C. Samples were sonicated using a Vibracell 75186 and a miniprobe 2 mm (Amp 80% // Pulse 10 off 0.8, 3 cycles). Proteins were then alkylated with 20 mM iodoacetamide for 30 min at room temperature in the dark. Subsequently, LysC (Promega) was added for the first digestion step (protein to Lys-C ratio = 80:1) for 3h at 30°C. Then samples were diluted down to 1 M urea with 100 mM Tris pH 7.5, and trypsin (Promega) was added to the sample at a ratio of 50:1 for 16h at 37°C. Proteolysis was stopped by adding Formic acid (FA) to a final concentration of 1 % (vol/vol). Resulting peptides were desalted using Sep-Pak SPE cartridge (Waters) according to manufactures instructions.

### LC-MS/MS of tryptic digest

LC-MS/SM analysis of trypsin-digested proteins (peptides) was performed on an Orbitrap Q Exactive Plus mass spectrometer (Thermo Fisher Scientific, Bremen) coupled to an EASY-nLC 1200 (Thermo Fisher Scientific). A home-made column was used for peptide separation [C_18_ 40 cm capillary column picotip silica emitter tip (75 μm diameter filled with 1.9 μm Reprosil-Pur Basic C_18_-HD resin, (Dr. Maisch GmbH, Ammerbuch-Entringen, Germany)]. It was equilibrated and peptide was loaded in solvent A (0.1 % FA) at 900 bars. Peptides were separated at 250 nL.min^-1^. Peptides were eluted using a gradient of solvent B (ACN, 0.1% FA) from 3% to 22 % in 160 min, 22% to 50% in 70 min, 50% to 90% in 5 min (total length of the chromatographic run was 250 min including high ACN level step and column regeneration). Mass spectra were acquired in data-dependent acquisition mode with the XCalibur 2.2 software (Thermo Fisher Scientific, Bremen) with automatic switching between MS and MS/MS scans using a top-10 method. MS spectra were acquired at a resolution of 70000 (at *m/z* 400) with a target value of 3 × 10^6^ ions. The scan range was limited from 300 to 1700 *m/z*. Peptide fragmentation was performed using higher-energy collision dissociation (HCD) with the energy set at 27 NCE. Intensity threshold for ions selection was set at 1 × 10^6^ ions with charge exclusion of z = 1 and z > 7. The MS/MS spectra were acquired at a resolution of 17500 (at *m/z* 400). Isolation window was set at 1.6 Th. Dynamic exclusion was employed within 45 s.

### Data processing

Data were searched using MaxQuant (version 1.5.3.8 and 1.6.6.0) (Cox and Mann 2008, Tyanova et al. 2016) using the Andromeda search engine (Cox et al. 2011) against home-made databases. The following databases were used. For *C. neoformans* KN99α, *C. deneoformans* JEC21 and *C. deuterogattii* R265 we used the recently updated proteomes (Wallace et al. 2020, Gröhs Ferrareze et al. 2021). The following search parameters were applied: carbamidomethylation of cysteines was set as a fixed modification, oxidation of methionine and protein N-terminal acetylation were set as variable modifications. The mass tolerances in MS and MS/MS were set to 5 ppm and 20 ppm respectively. Maximum peptide charge was set to 7 and 7 amino acids were required as minimum peptide length. A false discovery rate of 1% was set up for both protein and peptide levels. The iBAQ intensity was used to estimate the protein abundance within a sample (Schwanhäusser et al. 2011).

### Statistical analysis

All statistical analyses were performed using GraphPad Prism 8 software (GraphPad Software Inc.). Data sets were tested for normal distribution using Shapiro-Wilk or Kolmogorov-Smirnov normality tests. In the cases in which the data passed the normality test, they were further analyzed using the unpaired Student’s t test or ordinary one-way ANOVA. When at least one data set was nonnormally distributed, we used the nonparametric Kolmogorov-Smirnov or Kruskal-Wallis test. For the comparison of the survival curves, we used the Logrank (Mantel-Cox) test.

## 3. Results

### Diversity of cryptococcal EVs

Several groups have performed morphological studies of fungal EVs by electron microscopy (Rodrigues et al. 2007, Oliveira et al. 2009, Rayner et al. 2017, Bleackley et al. 2020). However, most of these studies used sample fixation-dehydration procedures for transmission electron microscopy (TEM), which can often affect the size and morphology of EVs (Van Der Pol et al. 2010, Chiang and Chen 2019). Cryo-EM imaging on rapidly-frozen samples at low temperature could potentially reduce sample damaging and artifacts caused by the addition of heavy metals, dehydration, or fixation steps (Orlov et al. 2017, Chiang and Chen 2019). Indeed, diverse morphologies of EVs derived from even a single mammalian cell type have been clearly revealed under cryo-EM (Zabeo et al. 2017). We therefore used cryo-EM and cryo-ET to analyze EVs purified from *C. neoformans*, in their near-native state.

Based on the optimized version of the EV purification protocol recently described by Reis and collaborators (Reis et al. 2019), we isolated EVs from *C. neoformans* reference strain KN99α, cultured on synthetic dextrose solid medium for 24h, in order to limit the carryover of potential contaminants. Cryo-ET tomograms allowed us to analyze 533 single vesicles, which were characterized according to their morphological aspects in regular (round-bilayered vesicles) and irregular (not rounded, bi- or multilayered vesicles) categories. Although a large proportion (81.4%) of the observed EVs had the typical round shape, 18.6% corresponded to irregular morphologies. Among them, we observed examples of multilayered vesicles, long tubular, flat, short tubular and miscellaneous morphologies **(Fig. S1; Table S2)**. However, it remains to be determined whether EVs with irregular morphologies are produced biologically or they appear as a consequence of the purification method.

Cryo-EM analysis showed a considerable polymorphism of EVs, with the two leaflets of the typical vesicular membrane readily visible for all EVs observed, and a few unstructured aggregates, thus confirming the quality of our preparation **(Fig. 1A)**. In *C. neoformans*, among the regular vesicles, only 10.8% appeared to have a smooth surface **(Fig. 1B and 1C)**; the majority of regular EVs (89.2%) were decorated with a fibrillar structure anchored to the lipid bilayer **(Fig. 1D and 1E).** Strikingly, regardless of the morphology, the majority of EVs analyzed (88.6%) appeared to be coated with this fibrillar material. We used cryo-ET to prepare a three-dimensional surface model of the EVs, using IMOD (Mastronarde and Held 2017) and UCSF Chimera (Pettersen et al. 2004) to further visualize their structure and fibrillar decoration **(Fig. 1F to 1H)**.

**Figure 1:**
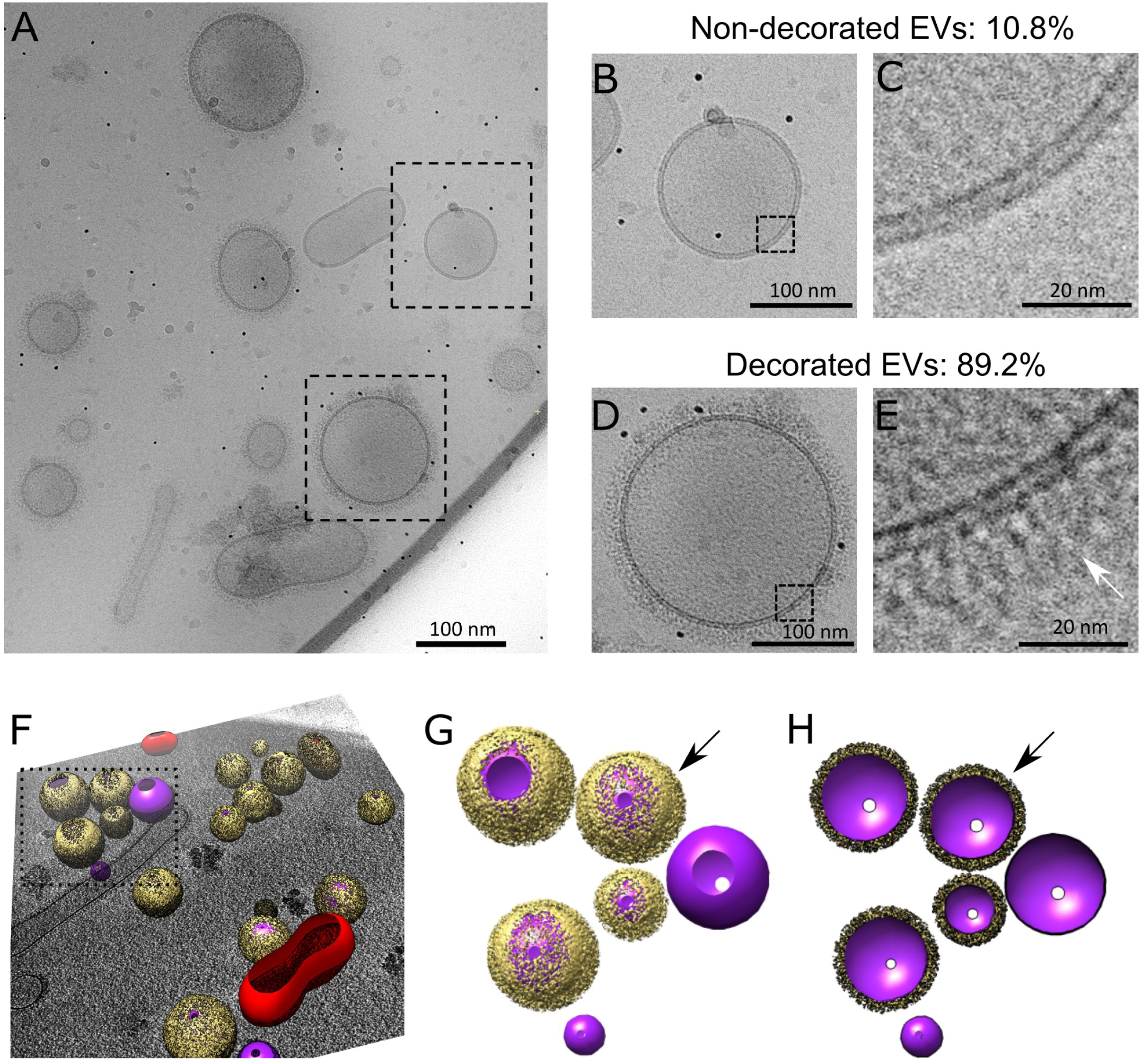
Cryo-electron microscopy analysis of *C. neoformans* extracellular vesicles (EVs). Cryo-EM analysis revealed a heterogeneous population of vesicles with diverse structural aspects, previously unappreciated in fungal EVs (A). As shown, the EVs were delimitated by a lipid bilayer (B to E), which showed either no decoration (in 10.8% vesicles, panels B and C) or a fibrillar decoration (arrows) in 89.2% of the EVs analyzed (panels D and E). Three-dimensional organization of the fibrillar decoration (yellow) on the membrane (purple) of EVs as revealed by cryo-electron tomography analysis (F), magnified in panels G and H. Full surface representation models as seen from top view (G). Same models clipped with clipping plane oriented perpendicular to line of sight (H). Data presented in this figure have been generated using images obtained using a Titan Krios (Thermo Scientific) transmission electron microscope.

Additional aspects of *C. neoformans* EV diversity, such as the distribution of size and decoration, were analyzed. NTA analyses showed a diameter size distribution from 80 to 500 nm and revealed a major peak of vesicle detection in the 150-nm-diameter range **(Fig. 2A),** in line with previous findings (Reis et al. 2019). We also analyzed the EV diameter frequency distribution by cryo-EM from 434-single regular EV captures **(Fig. 2B)**. The size distribution of vesicles tracked with NTA was different from the distribution of vesicles observed with cryo-EM, which revealed a wider range of EV diameter size, ranging from as small as 10 nm to 500 nm **(Fig. 2C; Video S1)**. Notably, smaller vesicles (< 100 nm) comprised a higher proportion of vesicles captured by cryo-EM than by NTA. Although cryo-EM has some statistical limitations, it nonetheless confirms the known bias of NTA towards larger EVs (Bachurski et al. 2019).

**Figure 2:**
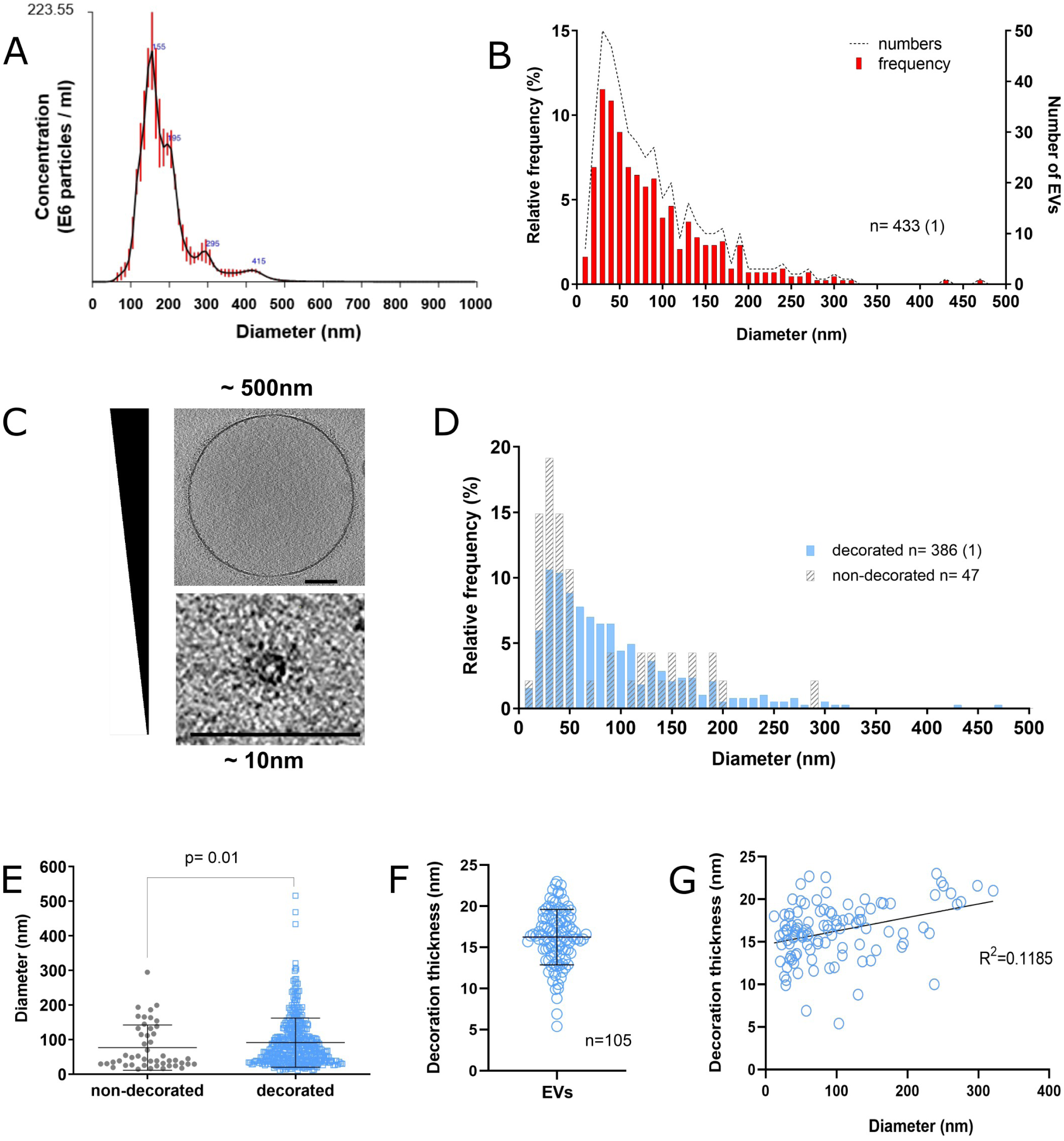
Analysis of size and structural diversity of *C. neoformans* EVs. NTA analysis of purified EVs revealed a size diameter ranging from 80 to 500 nm, with the highest distribution around 150 nm (A). Frequency distribution of EV diameters determined by CryoEM, a total of 434 regular EVs were analyzed. The analysis based on CryoEM tomograms revealed a wider range of EV size distribution, from 10 to 500 nm diameter, with the highest relative frequency below 100 nm (B). Cryo-EM images exemplifying EV size range. Scale bars corresponding to 100 nm (C). EV size distribution according to the presence or absence of surface decoration (D). Non-decorated EVs have a smaller diameter size distribution compared to decorated ones (E). Analysis of decoration thickness from Cryo-EM images from 105 single EVs (F). Analysis of a potential relationship between decoration thickness and EV diameter by linear regression (G). Data presented in this figure have been generated using images obtained using a Titan Krios (Thermo Scientific) transmission electron microscope. Error bars show means ± SD. Sample size (n) is indicated and, in brackets, the number of vesicles in that category that exceeded 500 nm in size.

Analysis of the EV size according to the presence or absence of the surface decoration revealed a different frequency distribution **(Fig. 2D)**, with non**-**decorated EVs showing a significantly smaller size distribution (*p* = 0.01, using nonparametric Kolmogorov-Smirnov test) compared to the decorated ones **(Fig. 2E).** Additionally, the analysis of the vesicular decoration in 105 single regular EVs revealed heterogeneity in their thickness, ranging from 5 to 23 nm with the average value close to 16 nm **(Fig. 2F)**. There was no correlation between vesicular diameter size and decoration thickness, as indicated by linear regression analysis **(Fig. 2G).** Therefore, the presence or absence of decoration, and even its thickness, does not depend on the size and shape of the EVs, revealing a previously unknown aspect of fungal EV diversity.

We analyzed EVs from two other pathogenic species of *Cryptococcus*, *C. deneoformans* strain JEC21 and *C. deuterogattii* strain R265. As expected, cryo-EM revealed a similar structure of the EV population in the three *Cryptococcus* species, the majority of EVs being decorated in *C. deneoformans* (72.4 %) and *C. deuterogattii* (81.4 %) **(Table S2)**. In contrast, *C. deuterogattii* EVs appeared to be smaller (median size = 48 nm) than those of *C. neoformans* (median size = 67 nm) and *C. deneoformans* (median size = 70 nm) **(Fig. 3A)**. In addition, the thickness of decoration is smaller in *C. deneoformans* and *C. deuterogattii* than in *C. neoformans* **(Fig. 3B),** suggesting a tight genetic control of these EV structural properties **(Fig. 3C, Fig. S2)**.

**Figure 3:**
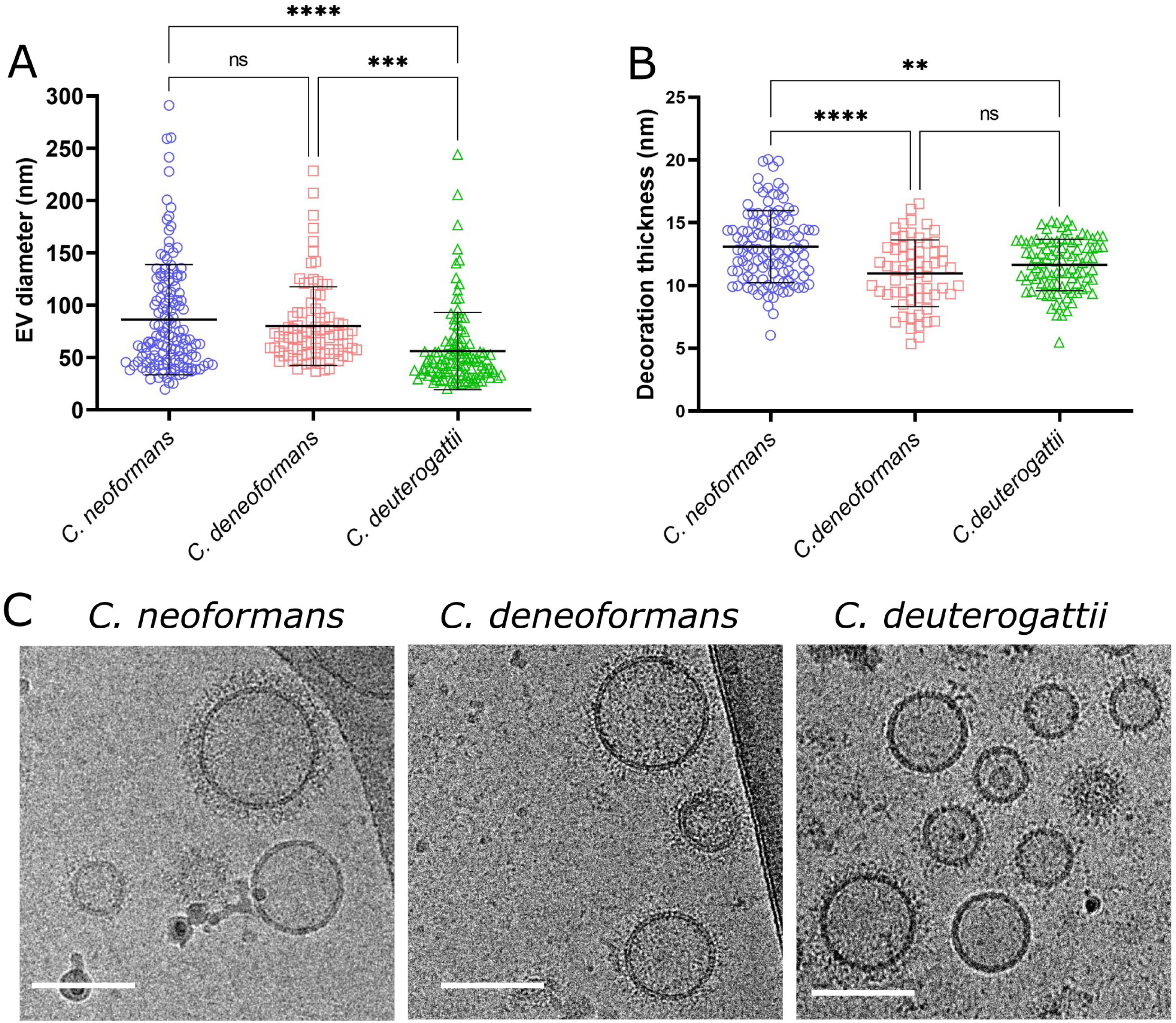
Comparative analysis of size and structural diversity of EVs in *C. neoformans*, C. deneoformans *and* C. deuterogattii. Analysis of EV diameters revealed a smaller size distribution in *C. deuterogattii* strain R265 and *C. deneoformans* strain JEC21 than in *C. neoformans* KN99α. The total numbers of vesicles analyzed were *C. neoformans* (n=143 for size and n=112 for decoration), *C. deneoformans* (n= 90 for size and n=63 for decoration), *C. deuterogattii* (n= 115 for size and n=95 for decoration) (A). Analysis of the decoration thickness revealed a smaller distribution for *C. deneoformans* and *C. deuterogattii* compared with *C. neoformans* (B). Illustrative images of size and decoration of EVs obtained from the three species. The data presented in this figure have been generated using images obtained using a TECNAI F20 transmission electron microscope (C). Error bars show means ± SD. Scale bars represent 100 nm.

### Cryptococcus EVs structural analysis

*C. neoformans* is an encapsulated microorganism, and its capsule is mostly composed of the polysaccharide glucuronoxylomannan (GXM), a critical virulence factor of this pathogenic yeast (O’Meara and Alspaugh 2012). GXM has been previously shown to be exported by EVs (Rodrigues et al. 2007). Therefore, we reasoned that the fibrillar decoration observed around the vesicles could be composed of GXM. We thus incubated *C. neoformans* EVs with the Alexa 488 labelled anti-GXM monoclonal antibody 18B7 (Casadevall et al. 1992), and analyzed the EV suspension by flow cytometry. More than 70% of the EVs obtained from the wild-type strain were recognized by this antibody **(Fig. 4A),** suggesting that most *C. neoformans* EVs are covered to some extent with GXM or derivatives thereof. While, EVs obtained from the acapsular mutant strain (*cap59Δ*) (Moyrand et al. 2007) showed negligible labelling (2.33%), following the same experimental approach **(Fig. 4B)**. Nevertheless, cryo-EM observation of *cap59Δ* EVs revealed similar fibrils as observed in the wild type EVs **(Fig. 4B)**. Moreover, cryo-EM analysis of EVs purified from *cap59Δ* suggested a similar percentage of decorated EVs (91.6%). Overall, these data suggest that, even though GXM covers most *C. neoformans* EVs, the visible fibrillar structures around them are not GXM-based. Cryo-EM analysis of EVs obtained from *C. albicans* SC5314 and *S. cerevisiae* S288C grown on SD medium showed a similar fibrillar decoration observed around *Cryptococcus* EVs **(Fig. 4C)**, reinforcing the notion that this structure is not GXM-based, since neither of these two yeasts can synthesize this capsular polysaccharide.

**Figure 4:**
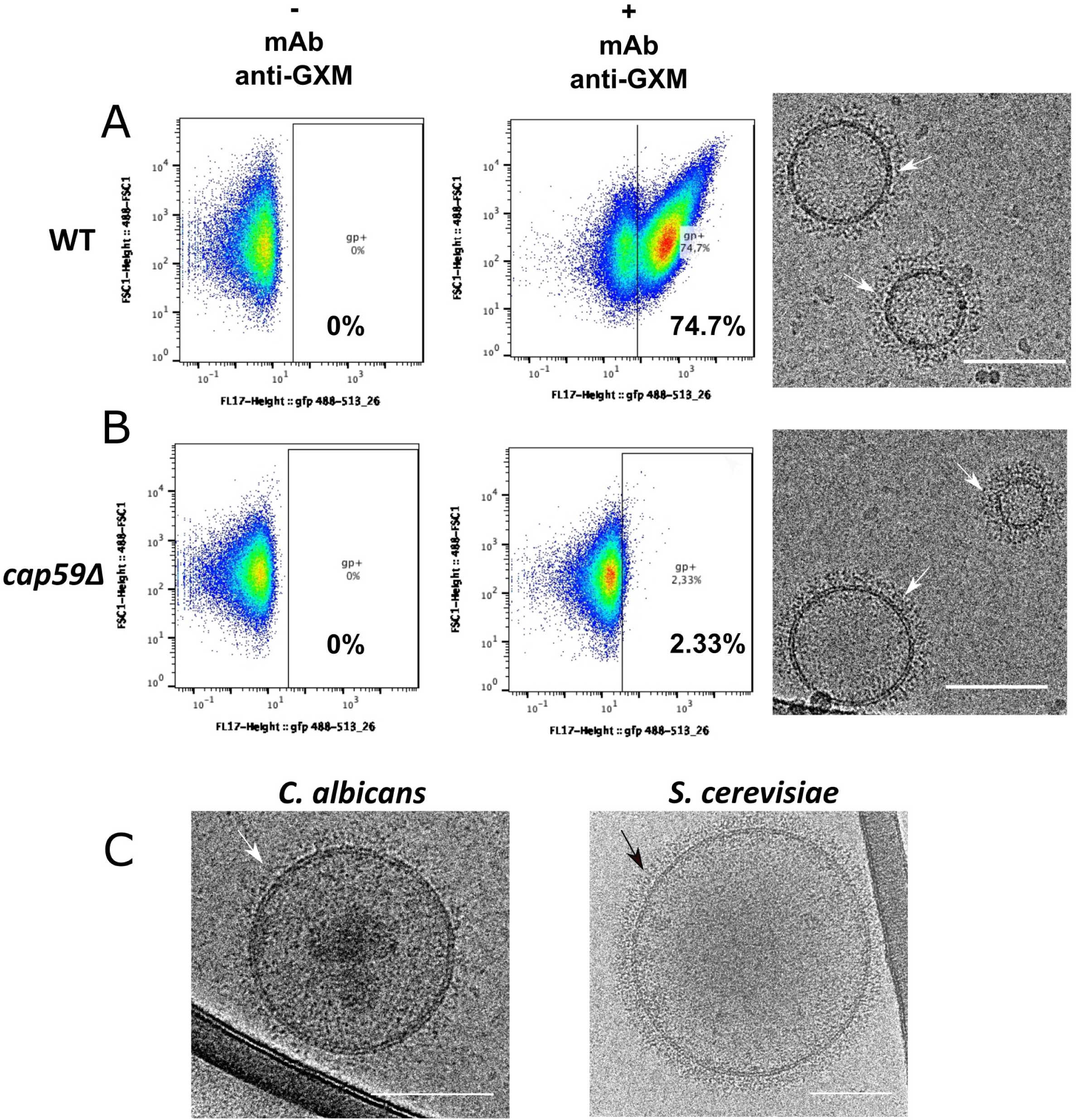
Flow cytometry analysis of *C. neoformans* EVs incubated with monoclonal anti-GXM antibody. FACS analysis of wild type (WT) and the acapsular *cap59Δ* EVs in PBS or in the presence of the monoclonal antibody raised against the capsular polysaccharide 18b7 (+ mAb anti-GXM) (A). The analysis revealed strong labelling of WT vesicles (74.7%), compared to the weak labelling in the mutant (2.33%), (B). Despite the important labelling difference, *C. neoformans* WT and *cap59Δ* strains released EVs bearing similar surface decoration, shown by the cryo-EM (arrows), as well as EVs obtained from other fungal species such as *C. albicans* and *S. cerevisiae* (C). These cryo-EM data have been generated using a TECNAI F20 transmission electron microscope. Scale bar represents 100 nm. This experiment was repeated twice with similar results.

We then reasoned that EV decoration could be protein-based and therefore performed proteomic analyses to further explore this novel fungal vesicular feature. Two proteomic analyses of *C. neoformans* EVs have been reported previously (Rodrigues et al. 2008, Wolf et al. 2014) wherein the authors identified 92 and 202 proteins associated with EVs in *C. neoformans*, respectively. However, neither quantitative nor enrichment of EV-associated proteins was performed in these two studies. Therefore, we performed EV proteomic characterization, together with an enrichment analysis in order to distinguish the proteins associated with EVs, from those related to potential carry-over aggregates, inevitably contaminating EV preparations.

In fungi, and more specifically in *Cryptococcus*, the relationship between RNA and protein abundances has been reported as nearly linear, due to the relatively minor contribution of posttranscriptional regulations to protein abundance (Wallace et al. 2020). We thus used cellular RNA abundance at 30°C, exponential phase (Wallace et al. 2020), as a proxy for cellular protein abundance, and for normalization of EV proteome data. *C. neoformans* EVs proteomic analysis was performed in experimental triplicate that produced a common list containing 1847 proteins **(Table S3).** Proteins were ranked according to their prevalence in the sample evaluating the average intensity-based absolute quantification (IBAQ) value of the three replicates. We then used the gene expression level as evaluated by RNA-seq analysis to calculate an enrichment coefficient comparing the expected value in the cells with the observed one in EVs. We thus identified 39 non-ribosomal proteins which were present both within the 100 most prevalent EV proteins overall and the 100 most enriched proteins **(Table S4)**. We considered these proteins as EV-associated proteins. Only 9 out of these 39 proteins were reported in previous proteomic analysis, emphasizing the necessity for proteomic data enrichment analysis. Of note, our study and those published before used different culture media, and distinct protocols of EV isolation, which might also explain the differences in protein composition that were presently observed.

To further explore how conserved the EV protein cargo across *Cryptococcus* species is, we proceeded with the same strategy to characterize the EV-associated proteins in two other cryptococcal species, *C. deneoformans* (strain JEC21) and *C. deuterogattii* (strain R265). We identified 38 and 48 EV-associated proteins for *C. deneoformans* and *C. deuterogattii*, respectively **(Table S3; Table S4)**. Overall, 71 EV-associated proteins were identified, 37 in at least two species, and 17 shared by all the three species **(Fig. 5A and B)**, supporting a conserved profile of the EV-associated proteins across *Cryptococcus* species, and the robustness of our analyses. Several families of proteins appeared to be typical of *Cryptococcus* EVs. The major one was the Chitin deactelylase Cda family (Baker et al. 2011), composed of three members present among the 17 EV-associated proteins identified in all three *Cryptococcus* species analyzed. Some other families like the putative glyoxal oxidase (Gox proteins), or the Ricin-type beta-trefoil lectin domain-containing protein (Ril), have one member common to all three species EVs (i.e. Gox2 and Ril1) whereas the other members are found in only two species (Ril2 and Ril3) or are specific of one species EVs (Gox1 and Gox3) **(Fig. 5C)**. We also identified three tetraspanin membrane proteins containing a SUR7/PalI family motif. Tsh1 and Tsh2 shared 32% identity in their amino acid sequence. Tsh1 is present in both *C. neoformans* and *C. deneoformans* EVs whilst Tsh2 was identified in both *C. neoformans* and *C. deuterogattii*. The third Sur7/PalI protein shares very little sequence homology beyond the SUR7 motif and is exclusive to *C. deuterogattii*. Two Sur7 proteins have been recently identified in *C. albicans* EVs, suggesting that they might represent a common EV marker present in fungal EVs (Dawson et al. 2020). Finally, two members of the previously described pr4/barwin domain Blp protein family (Chun et al. 2011) were present in *C. neoformans* and *C. deuterogattii* EVs but not in *C. deneoformans*. Similarly, the two ferroxidase Cfo proteins (Jung et al. 2008) were shown to be associated only with the *C. deuterogattii* EVs but not in the two other species.

**Figure 5:**
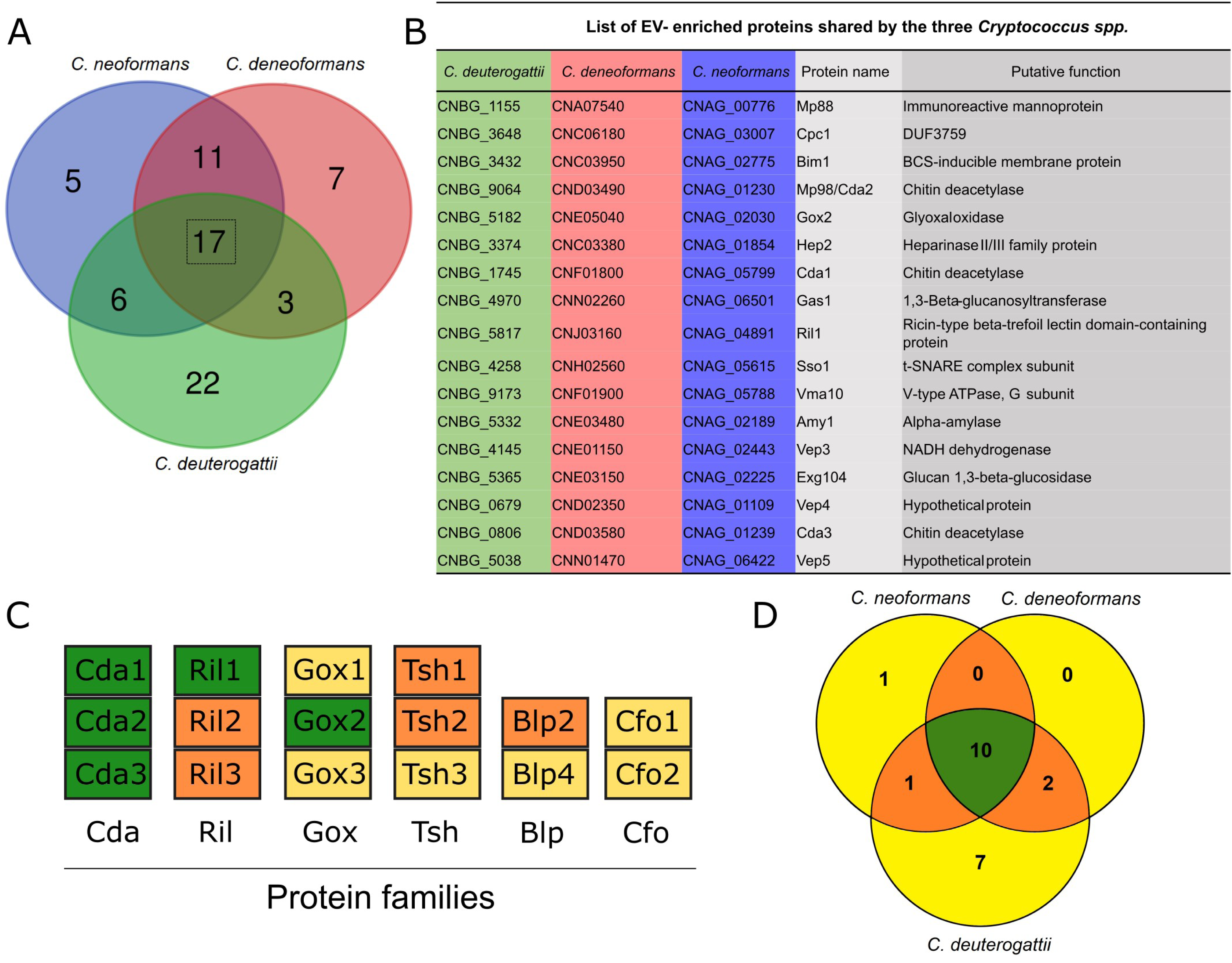
Analysis of *Cryptococcus spp* protein cargo. Venn diagram revealing shared and unique EV-associated proteins in *C. neoformans*, *C. deneoformans*, and *C. deuterogattii*. Seventeen proteins were identified to be associated with EVs in all three *Cryptococcus* species (A). List of the gene loci and the corresponding proteins commonly found in EVs released by the three species, which could be considered as putative cryptococcal EV-protein markers (B). Most of the proteins are predicted to be either GPI-anchored proteins, to contain a signal peptide or to possess other membrane domains, according to preGPI, signalP and TMHMM website, respectively. Six protein families appeared to be typical of *Cryptococcus* EVs, including the Chitin deacetylase family (Cda), the Ricin-type beta-trefoil lectin domain-containing protein family (Ril), the putative glyoxal oxidase family (Gox), the tetraspanin membrane proteins containing a SUR7/PalI family motif (Tsh), the pr4/barwin domain protein family (Blp), and the multicopper oxidase (Cfo). Among these families, the proteins present in all three species are shown in green, proteins present in two species in orange and proteins present in only one species in yellow (C). We also identify 21 putative GPI-anchored proteins, as predicted by PredGPI, and 10 of them were present in all three species (D).

Several enzymes associated with polysaccharide degradation and modifications were present in *Cryptococcus* EVs. Some of these proteins are specific to one species but others are present in two or all three EV proteomes. For instance, identification within the *Cryptococcus* EV core proteins of Gas1 (a 1,3-beta-glucanosyltransferase), Amy1 (an alpha amylase), Exg104 (a glucan 1,3-beta-glucosidase), Hep2 (a putative heparinase) together with the Gox, Cda and Ril proteins suggest functions of EVs in cell wall processes, as previously hypothesized in *S. cerevisiae* (Zhao et al. 2019). We also identified the BCS-inducible membrane protein (Bim1), recently described as a critical factor for cupper acquisition in *C. neoformans* meningitis (Garcia-Santamarina et al. 2020). Finally, several of the EV proteins identified here have no predicted function; we therefore named them Vep (Vesicles enriched protein). Bioinformatics analysis of the 71 EV-associated protein sequences suggested that 80% might be membrane-bound, 36 of them bearing at least one putative transmembrane domain as predicted by SignalP-5.0 (Almagro Armenteros et al. 2019) and/or TMHMM v. 2.0 (Krogh et al. 2001), and 21 being putative GPI-anchored proteins as predicted by PredGPI (Pierleoni et al. 2008), which is in good agreement with putative protein-based decoration. Reflecting the general specificities of these three proteomes, the GPI-anchor EV-proteomes of *C. neoformans* and *C. deneoformans* are nearly identical, whereas *C. deuterogatii* is more diverse (**Fig. 5D**).

Mature GPI-anchored proteins can also be membrane-bound and are predicted to be highly mannosylated in *Cryptococcus* and other fungi (Levitz et al. 2001, de Groot et al. 2003). We thus reasoned that these mannosylated proteins might represent the EV decorations observed by cryo-EM. To test this hypothesis, we incubated EVs with ConA conjugated to Alexa Fluor 488, and further analyzed by flow cytometry. Our results demonstrated that over 98.5% of vesicles were recognized by this lectin, confirming the presence of mannosylated proteins on the EV surface **(Fig. 6)**. Similarly, EVs obtained from acapsular *cap59Δ* mutant strain also showed a high percentage of staining (95.5%). Accordingly, EV treatment with proteinase K was associated with a nearly complete loss of ConA labelling of both WT and *cap59Δ* EVs (**Fig. 7**), overall suggesting that the outer vesicle decoration may be composed primarily of mannoproteins.

**Figure 6.**
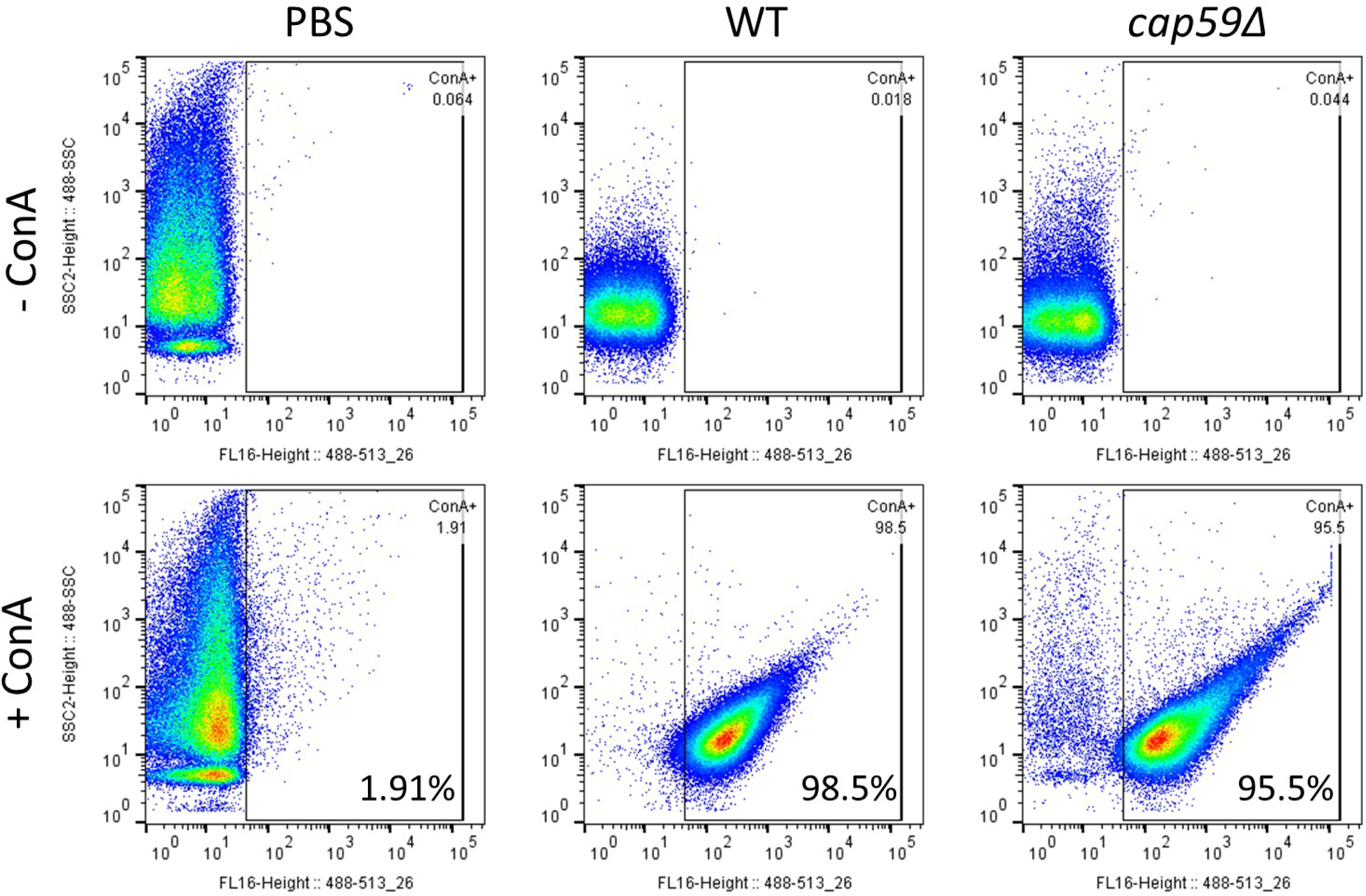
Flow cytometry analysis of *C. neoformans* EVs incubated with GFP-labelled ConA. FACS analysis of EVs obtained from *C. neoformans* wild type and *cap59Δ* cells. EVs were incubated with ConA-Alexa Fluor 488 conjugated lectin. After ultracentrifuge washing, the EV pellets were mixed in BD Trucount tubes (BD Biosciences), containing a known number of fluorescent beads as internal control. The number of events for each reading was fixed to 100,000 events and the percentage and intensity of ConA labeling were recorded. This experiment was repeated three times with similar results.

**Figure 7.**
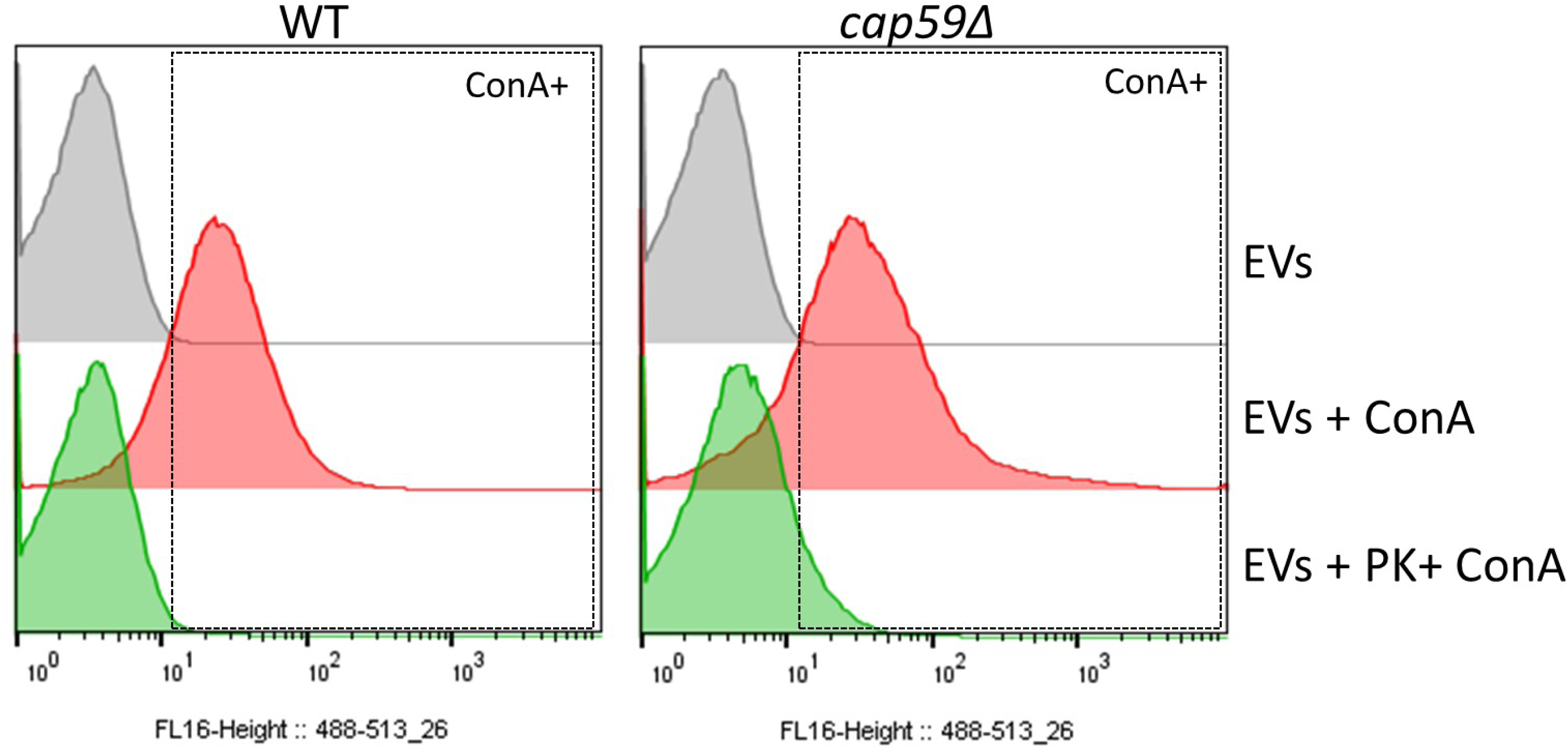
EV proteinase K treatment reduces ConA binding. FACS analysis of EVs obtained from *C. neoformans* WT and *cap59Δ* cells after proteinase K treatment. Proteinase K-treated EVs were submitted to ConA labelling, ultracentrifuge washed and analyzed by flow cytometry. EV pellets were mixed in BD Trucount tubes (BD Biosciences), containing a known number of fluorescent beads as an internal control. The number of events for each reading was fixed to 100,000 events and the percentage and intensity of ConA labeling were recorded. EVs treated using the same protocol but omitting the enzyme were used as controls.

Several genes have been implicated in protein glycosylation in *C. neoformans*. For instance, *ALG3* encodes a dolichyl-phosphate-mannose-dependent α-1,3-mannosyltransferase, deletion of which is associated with the production of truncated protein-associated neutral *N*-glycans and a reduction in virulence (Thak et al. 2020). Similarly, *KTR3* and *HOC3* encodes α1,2-mannosyltransferase and α1,6-mannosyltransferase, respectively, regulating *O*-glycan structure and pathogenicity of *C. neoformans* (Lee et al. 2015). We reasoned that the deletion of some of these genes could alter EV production and structure. We first analyzed EV production in *alg3Δ*, *hoc3Δ*, and *ktr3Δ* strains by evaluating the quantity of sterol in our EV preparations. We did not observe any significant alternation in EV production nor in the percentage of ConA positive EVs in any of these deletion mutants (**Fig. 8A, 8B**). Nevertheless, the percentage of *alg3Δ* EVs labelled by ConA was slightly reduced **(Fig. 8B)** and *alg3Δ* EV decorations were less thick than wild type EVs, as revealed by cryo-EM observation (**Fig. 8C, 8D; Table S2**).

**Figure 8.**
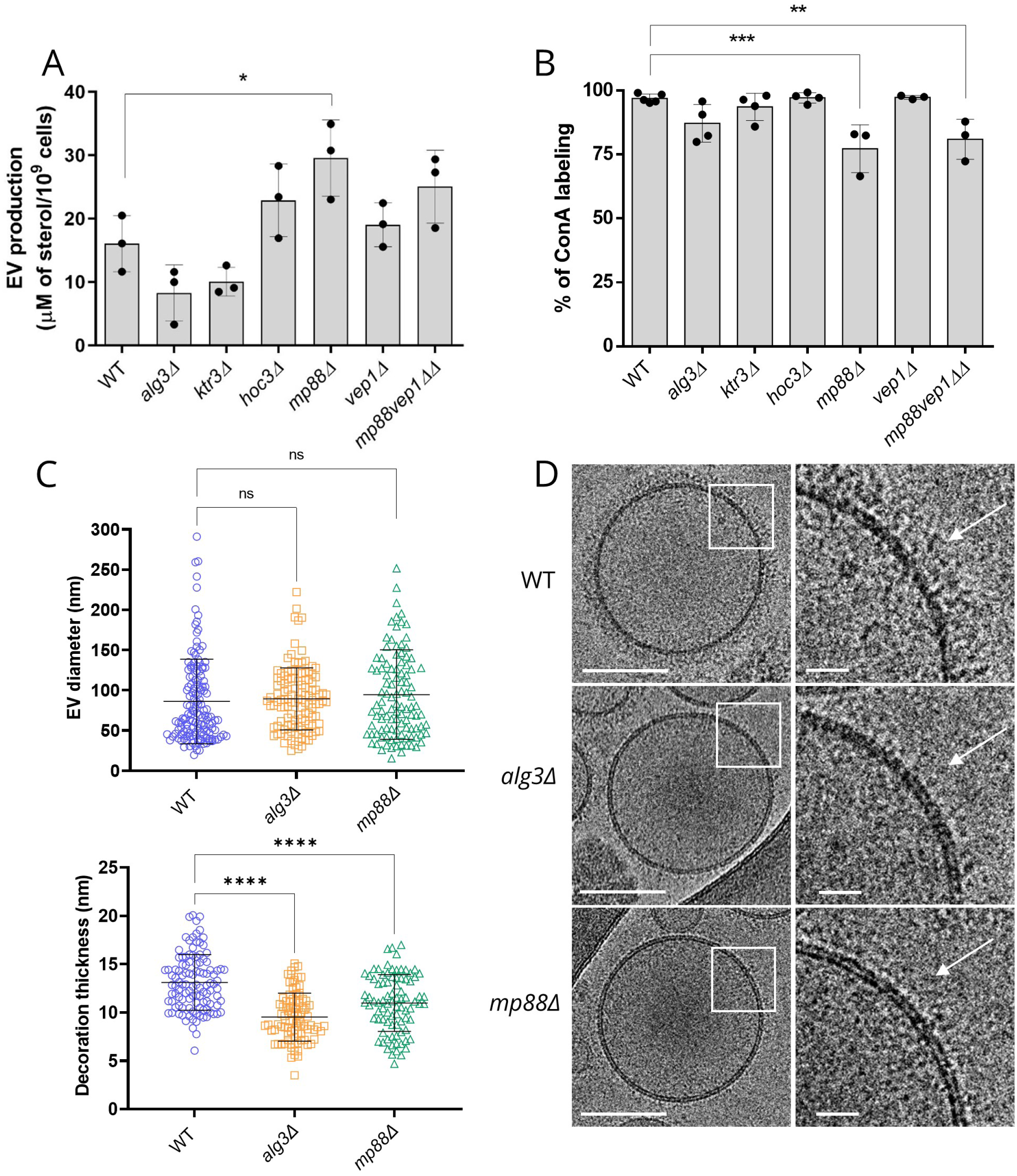
Analysis of *C. neoformans* mutant strain EVs. Evaluation of EV production by the different mutant strains as estimated by the measure of the sterol concentration using the Amplex™ Red Cholesterol Assay Kit (A). Impact of the different mutations on the percentage of ConA-labelled EVs as estimated through flow cytometry (B). Analysis of EV size diameter in the *mp88Δ* and *alg3Δ* mutant strains as compared to the wild type (WT). The total number of vesicles analyzed were WT (n = 143 for size and n = 112 for decoration), *mp88Δ* (n = 107 for size and n = 86 for decoration), *alg3Δ* (n = 119 for size and n = 92 for decoration) (C). Analysis of the decoration thickness revealed a smaller distribution associated with *ALG3* or *MP88* deletions, as exemplified by illustrative images from the three strains (D). The cryo-EM images were obtained using a TECNAI F20 transmission electron microscope. ConA labelling and sterol measurements were done for at least three independent biological replicates Error bars are represented as means ± SD. Scale bars represent 100 nm in C and 20 nm in D. (E)

The two most abundant *C. neoformans* EV proteins, Mp88 and Vep1/CNAG_03223 are GPI-anchored and represent 23.7% of the total identified proteins. Mp88 is a basidiomycete specific protein originally identified as a major *C. neoformans* immunoreactive mannoprotein stimulating T cell responses (Huang et al. 2002). Vep1 (Vesicles Enriched Protein 1) is protein of unknown function sharing no homology with any *C. albicans* or *S. cerevisiae* protein. In all three *Cryptococcus* species, Mp88 (Huang et al. 2002) was the most prevalent EV protein. In *C. deuterogattii* EVs, in which the Vep1 protein is not present, Mp88 represents 35.4% of all EV proteins. We constructed the corresponding single and double mutant strains for *MP88* and *VEP1* and tested their EVs for ConA binding. These mutations did not strongly affect EV production although *MP88* deletion was associated with a slight increased production as compared to the wild type strain (**Fig. 8A**). However, both *mp88Δ* and *mp88Δ vep1Δ* EVs displayed a limited but statistically significant reduction of the ConA bound EVs as compared to EVs from wild-type strain (**Fig. 8B**). Accordingly, cryo-EM analysis of *mp88Δ* EVs revealed an associated reduction of the decoration thickness (**Fig.8C, 8D**) without any change in EV size distribution (**Fig. 8C; Table S2; Fig. S2**) suggesting that cryptococcal EVs might bear a highly complex decoration, probably formed from a dynamic combination of mannoproteins.Combining all these data, we propose a model for cryptococcal EV structure, in which, EVs are decorated by mannosylated proteins and covered by GXM **(Fig. 9)**.

**Figure 9:**
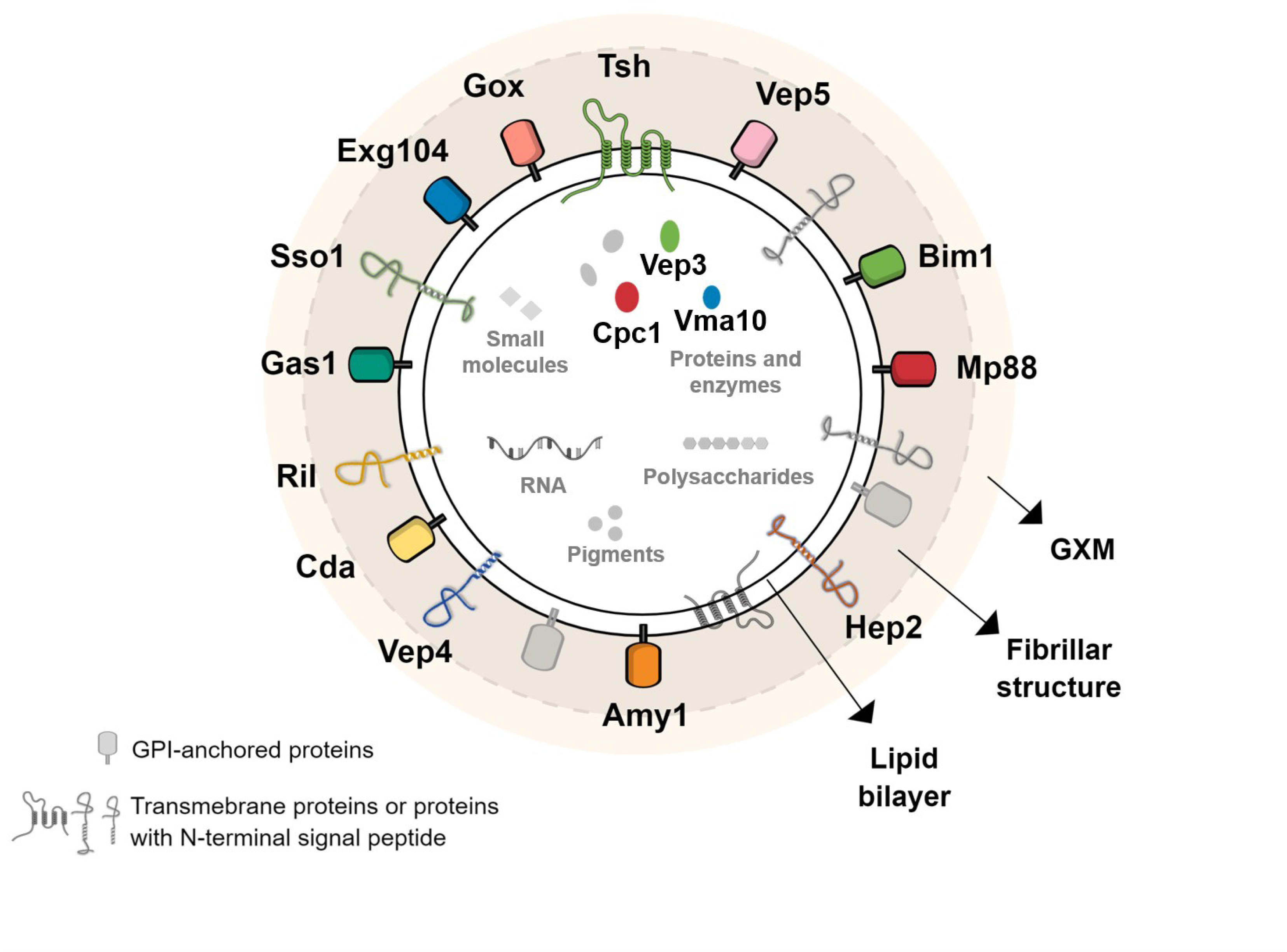
Model of simplified molecular structure and composition of *Cryptococcus* EVs. In accordance with previous reports and in the light of our data, a new model of *Cryptococcus* EVs is suggested, where the outer layer is composed of the capsular polysaccharide glucuronoxylomannan (GXM), and the lipid bilayer is covered by many proteins, including mannoproteins, making the visible fibrillar structure resolved by cryo-EM. Most of the proteins are predicted to be GPI-anchored, to contain a signal peptide or to possess other membrane domains, according to preGPI, signalP and TMHMM, respectively. Three proteins, the hypothetical protein Cpc1, the putative V-type ATPase (Vma10) and the Vep3 are predicted to be soluble. It is still unclear if these proteins are indeed inside the vesicular lumen or linked to another transmembrane protein. For simplification, the lipid content was not explored, but previous works shown the presence of sterol, phospholipoids and sphingolipids. Additionally, *Cryptococcus* EVs were also described to contain other cargoes, such as RNA, pigments, small molecules, and polysaccharides, including GXM, as detailed in plain text.

### EVs for immunization and protection against cryptococcal infection

Proteomic analysis of the *C. neoformans* EVs identified many immunogenic proteins, including Mp88, the members of Gox and Cda families and some Vep proteins previously tested as vaccine candidates against cryptococcosis (Specht et al. 2017, Hester et al. 2020). Moreover, some of these proteins were also found to be enriched in *C. deneoformans* and *C. deuterogattii* EVs (Mp88, Cda1, Cda2, Cda3, and Gox2), suggesting that secretion of these immunogenic molecules via EVs could be a conserved feature across different species. Taking into account that cryptococcal EVs have been shown to be immune modulators (Freitas et al. 2019) and may impact the pathophysiology of the infection (Bielska et al. 2018, Hai et al. 2020), we reasoned that EVs could be used for immunization against cryptococcosis, avoiding the need for recombinant protein purification and adjuvant usage. The usage of fungal EVs has been previously suggested as a promising vaccine strategy (Vargas et al. 2015, Colombo et al. 2019, Freitas et al. 2019, Vargas et al. 2020). However, to date cryptococcal EVs have not been tested in murine infection models.

In a pilot experiment, we obtained EVs from *C. neoformans* wild-type strain and the acapsular *cap59Δ* mutant, used them to immunize BALB/c mice in two different EV-protein dosages (1 and 10 μg) via intraperitoneal injections; control group was injected with only PBS (four mice in each group). After three immunizations, anti-EV-antibody response was evaluated in the mouse sera. Regardless of the EV origin, all the immunized mice produced antibodies against vesicular proteins, as revealed by Western Blot **(Fig. 10A)**. Forty days after the last immunization, mice were challenged intranasally with *C. neoformans* wild type strain (1 x 10^4^ yeasts per mouse), and their survival were monitored post-infection. All EV-immunized mice survived longer than the non-immunized ones and immunization with both doses of *cap59Δ* EVs statistically significantly prolonged the survival of the mice **(Fig. 10B)** To note, the total carbohydrate per 100 μg of EV-proteins were approximately 22 μg and 3 μg, respectively, for wild-type and *cap59Δ* mutant, as analyzed by gas-chromatography analyses (Richie et al. 2009). We then confirmed this result using a larger number of mice (10 mice per group). Since the highest dose of EVs from the acapsular mutant rendered the best protection, we decided to proceed only with EVs from *cap59Δ* strain (10 μg per mouse). After immunizations with EVs, the anti-EV-antibody response in the mice was analyzed; all immunized mice produced antibodies against vesicular molecules **(Fig. 10C)**. Following, the mice were challenged with *C. neoformans* wild type strain (1 x 10^4^ yeasts per mouse), and their survival was monitored post-infection. EV-immunization led to a significant prolonged survival (*p* = 0.0006) **(Fig. 10D)**, thus confirming the promising potential usage of EV-based protection against *Cryptococcus*.

**Figure 10.**
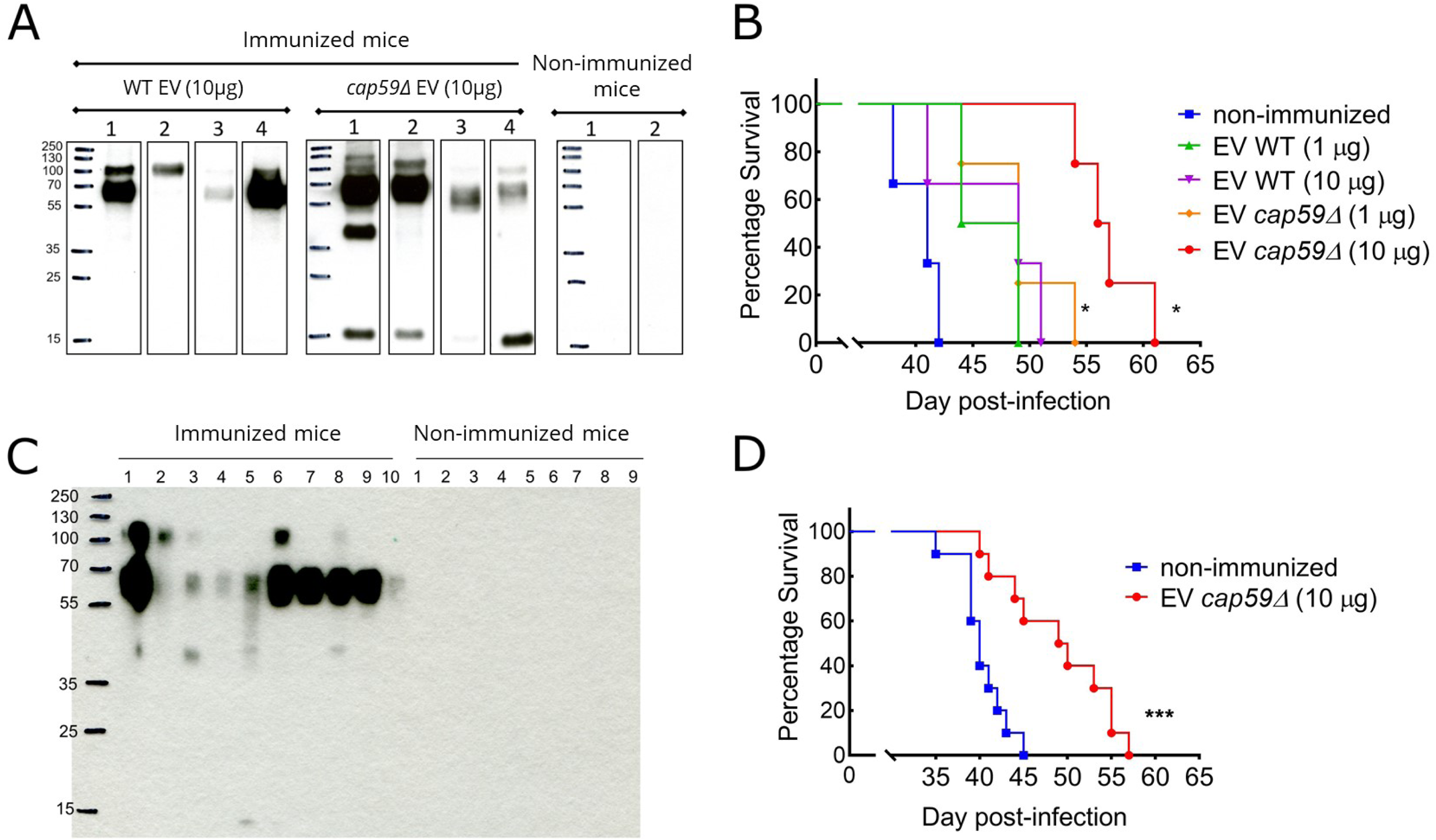
Vaccination assays using *C. neoformans* EVs. Female 6-weeks old BALB/c mice were immunized with *C. neoformans* EVs via intraperitoneal injection, followed by intranasal infection with 1 x 10^4^ yeasts of wild-type (WT) *C. neoformans*, and the mouse survival was monitored. In the first pilot experiment, mice (n = 4 per group) were immunized with EVs from wild type or *cap59Δ* strain (1 and 10 μg in 100 μL of PBS) and control mice were injected with 100 μL PBS. Western Blot using mouse sera against fungal EV confirmed that all immunized mice produced antibodies against EV proteins (A). All EV-immunized mice survived longer than the non-immunized ones, but the immunization with *cap59Δ* EVs rendered a significantly prolonged mouse survival (\**p* = 0.01) (B). For the second set of experiment, mice (n = 10 per group) were immunized with EVs from *cap59Δ* mutant strain (10 μg/100 μL in PBS) and control mice were injected with 100 μL PBS. Again, Western blot using mouse sera against fungal EVs confirmed that all immunized mice produced antibodies against EV proteins(C). EV-immunized mice showed significantly prolonged survival (\**p* = 0.0006) compared to the non-immunized group (D). Comparison of the survival curves was made by GraphPad Prism 8, using the Log-rank (Mantel-Cox) test.

## 4. Discussion

Studies on fungal EVs have gained much attention during recent years (Rizzo et al. 2020). Although data from both pathogenic and nonpathogenic species highlight their importance in diverse biological contexts, knowledge on fungal EVs is still limited, mostly due to their nanometer size and the technical hurdles intrinsic to the methods applied for their characterization (Rizzo et al. 2020). Here we used cutting edge technologies to revisit *Cryptococcus* EVs. Our cryo-EM analysis produced an unprecedented quality of EV images and resolved the fibrillar structure decoration as a new aspect on fungal EVs.

Our hypothesis is that EV decoration is not capsular polysaccharide GXM-based but mainly composed of mannoproteins. This is supported by two independent experiments. First, we demonstrated that although GXM most probably surrounds the vesicles, it is not necessary for the presence of decoration. Thus, EVs produced by an acapsular strain of *C. neoforman*s are not bound by a GXM specific antibody yet still display decoration. Secondly, *C. albicans* and *S. cerevisiae* EVs are also decorated, although none of these yeasts produced a capsular polysaccharide. Nonetheless, our study revealed that the deletion of single mannoproteins, such as the GPI-anchored proteins Mp88 and Vep1, was not sufficient to completely remove the EV decoration, suggesting that this structure has a highly complex and dynamic composition, including several mannoproteins.

Indeed, previous reports in *C. albicans* showed that the role of GPI-anchored proteins are redundant and single mutants mostly displayed minor phenotypes, if any (Plaine et al. 2008). Interestingly, Johansson and coworkers performed cryo-EM analysis of *Malassezia sympodialis* EVs, demonstrating no (Johansson et al. 2018) evident decoration on their surfaces (Johansson et al. 2018). Comparative genomic analysis suggested that this lipophilic pathogenic yeast, living on the skin (Theelen et al. 2018), lacks the N-glycosylation pathway and possesses only a very small number of GPI-anchor proteins (Gioti et al. 2013). Accordingly, *M. sympodialis* cells lack the extensive mannan outer fibrillar layer, which can be easily observed at the surface of the cell wall of most yeasts including *S. cerevisiae* or *C. albicans* (Gioti et al. 2013, Muszewska et al. 2017). Therefore, it is very tempting to hypothesize that this absence of mannans in *M. sympodialis* could explain the absence of EV decoration, supporting the idea that EV decoration in *Cryptococcus* species is mannoprotein-based. Previous proteomic analysis of fungal EVs identified putative mannoproteins, suggesting that this decoration is a common feature of fungal EVs (Bleackley et al. 2019, Dawson et al. 2020, Karkowska-Kuleta et al. 2020, Rizzo et al. 2020). Accordingly, flow cytometry experiments showed that *C. glabrata* EVs can be labelled by ConA (Karkowska-Kuleta et al. 2020). Putative fibril-like structures have also been reported at the surface of *Aspergillus fumigatus* EVs produced during cell wall regeneration (Rizzo et al. 2020).

In addition, we performed proteome analysis of EVs from *S. cerevisiae and C. albicans* grown in the same conditions as *Cryptococcus* species, and confirmed the presence of number of cell wall and GPI-anchored proteins in their EVs (Vargas et al. 2015, Zhao et al. 2019, Dawson et al. 2020). We also confirmed the presence of diverse antigenic proteins associated with EVs in *C. albicans,* reinforcing the notion that this feature might be a general aspect of pathogenic fungal EVs (**Table S5**). Whereas the presence of decoration seems to be a hallmark of fungal EV, it is not specific to this kingdom (Macedo-da-Silva et al. 2021). Although EVs bearing visible structures on their surface have not been commonly reported, a recent cryo-EM analysis of EVs derived from human breast cell lines overexpressing hyaluronan synthase 3-(HAS3) suggested the presence of fibril-like structures on their vesicle surface (Noble et al. 2020). Additionally, EVs from poliovirus-infected cells contain ‘protein structures with globular heads on a stalk’ around the membrane (Yang et al. 2020). Nevertheless, it is still unclear how often this feature is present among the whole EV population, and what the composition of these surface structures is.

Previous studies explored the size and morphology of fungal EVs, mostly by techniques such as electron microscopy (TEM, SEM), dynamic light scattering (DLS), and NTA (Albuquerque et al. 2008, Rodrigues et al. 2008, Wolf et al. 2014, Vargas et al. 2015, Wolf et al. 2015, Bielska and May 2019). Here we show that cryptococcal EVs are more heterogeneous than previously recognized in terms of size distribution and morphotypes. Our cryo-EM analysis revealed that the peak of EV size distribution was smaller than 100 nm, and substantially different from size distribution observed by NTA and from that previously found from *C. neoformans* and *C. deuterogattii* EVs using NTA and DLS approaches (100 to 300 nm) (Reis et al. 2019). Moreover, our study revealed not only the presence of regular EVs but also tubular, flat, and multilayered EVs. Although the different EV morphologies were previously identified in many fungal pathogens (Albuquerque et al. 2008, Rodrigues et al. 2008, Tefsen et al. 2014, Vargas et al. 2015), some vesicular shapes found in this work have not previously been reported. Thus, membrane tubule structures (memtubs) budding from the plasma membrane were found in the arbuscular fungus *Rhizophagus irregularis*, suggesting that different shapes of membranous structures could appear during fungal growth (Roth et al. 2019). Additionally, tubular and other morphologies were also found in EV populations obtained from human biological fluids (Arraud et al. 2014, Emelyanov et al. 2020). Although these data suggest that diverse structures could be part of the native EV population, the cellular origins of these structures are still unknown, and we cannot rule out the possibility of them being artifacts resulted from the filtration step of the commonly used EV isolation protocols.

In this study, we demonstrated that the three *Cryptococcus* species release both decorated and undecorated EVs, adding another previously unappreciated aspect to fungal EV diversity. As hypothesized before, this result also suggests the existence of at least two different pathways involved in EV biogenesis (Oliveira et al. 2010, Oliveira et al. 2013, Bielska and May 2019, Rizzo et al. 2020). It is, therefore, reasonable to hypothesize that decorated EVs could be shed from the fungal plasma membrane, “stealing” cell membrane proteins when they bud out. Interestingly, the decorated EVs have larger size distribution than the undecorated ones, in good agreement with what would be typical microvesicles in mammals. In this hypothesis, the enrichment of tetraspanin membrane proteins containing a SUR7/PalI family motif might indicate that decorated EVs could be specifically shed from the Sur7 specialized plasma membrane domains. This model could be extended to other fungi as

Sur7 proteins have been recently identified as EV-protein markers in *C. albicans* and in the wheat pathogen *Zymoseptoria tritici* (Dawson et al. 2020, Hill and Solomon 2020). This latter hypothesis, together with whether or not the smaller undecorated EVs are a result of the endosomal secretory pathways, thought to be exosomes being released by multivesicular bodies (MVB), still needs to be further explored. Interestingly, the characterization of decorated and undecorated EVs as microvesicles and exosomes, respectively, has previously been proposed (Noble et al. 2020). This hypothesis and our current results are supported by a recent study of *A. fumigatus* EVs in the absence of a cell wall. EVs were formed at the plasma membrane level and they contained a number of plasma membrane proteins (Rizzo et al. 2020).

Our work suggests that cryptococcal EV cargo contains proteins involved in diverse biological processes, including Mp88 and members of Cda and Gox families, which have been suggested as immunomodulators (Specht et al. 2017, Hester et al. 2020). Since the novel surface structure on fungal EVs resolved by cryo-EM resembles the spike complexes on viral envelopes (Neuman et al. 2006, Zanetti et al. 2006), we reasoned they may be useful as a vaccine platform. Numerous efforts are underway to develop vaccines against fungal infections, although none have yet been approved for human use (Nami et al. 2019). It was previously shown that the pre-treatment of *Galleria mellonella* larvae with fungal EVs stimulated a protective response against a lethal challenge with *C. albicans*, C*. neoformans* or *A. fumigatus* (Vargas et al. 2015, Colombo et al. 2019, Brauer et al. 2020). More recently, it was also demonstrated that *C. albicans* EVs were also able to elicit a protective effect against murine candidiasis (Vargas et al. 2020). Interestingly, *C. neoformans* EVs show immunoreactivity with sera from patients with cryptococcosis, indicating that EV-associated proteins are produced during cryptococcal infection (Rodrigues et al. 2008). Prophylactic immunization is one of the effective methods to prevent cryptococcal infection, and several cryptococcal antigens have been tested for their vaccination potential (Caballero Van Dyke and Wormley 2018, Ueno et al. 2020). However, the *in vivo* immunoregulatory role of EVs have largely remained unknown (Robbins and Morelli 2014).

In our study, antibody responses in cryptococcal EV-immunized mice indicate that the EVs can elicit an adaptive immune response in the absence of any adjuvants or carriers, unlike other antigenic proteins of *Cryptococcus* (Specht et al. 2017). It is also important to note that immunization using *C. neoformans* heat-killed cells does not elicit protection in a murine model of infection (Masso-Silva et al. 2018). EV-based vaccination data obtained by other groups using an invertebrate model suggest that innate immunity might also be involved (Vargas et al. 2015, Colombo et al. 2019). As *Cryptococcus* predominantly infects immunocompromised hosts, it will be worth checking the role of EVs in eliciting trained immunity, wherein innate immune cells develop memory-like response against an antigen upon repeated exposure (Hole et al. 2019, Mulder et al. 2019). The mechanisms, and the responsible immune cell types leading to prolonged survival in our murine infection model, remain to be deciphered. Although EV immunization was not sufficient to prevent death, we believe that adjusting the antigens exposed on EV surface could potentially increase the protective effect. In that sense, the fact that EVs from *C. neoformans* WT and the acapsular mutant did not lead to the same level of protection is an encouraging data.

Overall, the fantastic power of cryo-EM, together with several innovative analyses, has enabled us to draw a new model of fungal EVs and revealed new aspects of their diversity, suggesting different biosynthetic pathways. This model supports new strategies to construct vaccines against these too often neglected infectious diseases. It also opens the door to more questions concerning the origin and the fate of fungal EVs.

## Acknowledgements

This work was supported by a CAPES COFECUB grant n°39712ZK (to GJ, MLR, LA and LN). JR was supported by the CAPES-COFECUB Franco-Brazilian Research Exchange Program (88887.357947/2019-00) and by a Pasteur-Roux-Cantarini fellowship of Institut Pasteur. SSWW was supported by CEFIPRA/ANR-DFG-AfuINF grant and Pasteur-Roux-Cantarini postdoctoral fellowship. VA was supported by ANR-DFG AfuINF and Indo-French Centre for the Promotion of Advanced Research (CEFIPRA; Grant No5403-1) grants. Jean-Marie Winter from the NanoImaging Core at Institut Pasteur is acknowledged for his support image acquisition. The NanoImaging Core was created with the help of a grant from the French Government’s ‘Investissements d’Avenir’ program (EQUIPEX CACSICE – “Centre d’analyse de systèmes complexes dans les environnements complexes”, ANR-11-EQPX-0008). The Falcon II equipping the F20 microscope at the UBI used during this study was also financed by the Equipex CACSICE (grant number ANR-11-EQPX-0008). M.L.R. is supported by grants from the Brazilian Ministry of Health (grant 440015/2018-9), Conselho Nacional de Desenvolvimento Científico e Tecnológico (CNPq; grants 405520/2018-2 and 301304/2017-3), and Fiocruz (grants VPPCB-007-FIO-18 and VPPIS-001-FIO18)MLR, JR, and FCGR also acknowledge support from Coordenac☐ão de Aperfeic☐oamento de Pessoal de Nível Superior (CAPES, finance code 001) and the Instituto Nacional de Cie☐ncia e Tecnologia de Inovac☐ão em Doenc☐as de Populac☐ões Negligenciadas (INCT-IDPN). MLR is currently on leave of an associate professor position at the Instituto de Microbiologia Paulo de Góes of the Universidade Federal do Rio de Janeiro. We thank Adèle Trottier for her help in the preparation of EV samples

## Notes

### Competing Interest Statement

The authors have declared no competing interest.

### Summary of Updates

First, additional cryo-EM analyses of EVs isolated from two additional Cryptococcus species have been performed, confirming the diversity of fungal EVs and the presence of decoration in most of them. These experiments also revealed species specificities in EV size and decoration thickness. Second, our new experiments showed that proteinase K treatment strongly affect ConA labelling of EVs, supporting our model in which mannoproteins are at the surface of EVs. Third, we performed nano flow cytometry analyses and cryo-EM experiments to study N-and O-glycosylation mutant (alg3, ktr3, hoc3) EVs. These experiments revealed that ALG3 deletion is associated with reduced ConA labelling and decoration thickness. Fourth, we performed additional nano flow cytometry and cryo-EM analysis on an mp88Δ mutant strain revealing the impact of this mutation on EV ConA labelling and decoration thickness. Fifth, in this new version of the manuscript, we studied the proteomes of C. albicans and S. cerevisiae EVs showing that mannoproteins and GPI anchor proteins, including immunogenic ones, are common features of fungal EVs. Overall, all these data strongly support our previous conclusions and strengthen our new structural model of fungal EVs.

